# Attenuated adaptive growth of interpersonal synchrony in autism

**DOI:** 10.64898/2026.06.02.729457

**Authors:** Jinhwan Kwon, Hiromi Kotani

## Abstract

During social interactions, people continuously align their movements and rhythms, a process known as interpersonal synchrony that supports rapport, mutual understanding, and smooth communication. In autism spectrum disorder (ASD), previous studies have often reported atypical or reduced synchrony, but most have relied on aggregate or session-averaged measures that may miss how coordination develops over time. It therefore remains unclear whether interactional differences in autism reflect a general reduction in synchrony or altered temporal dynamics of interpersonal coordination. We examined the temporal dynamics of head-movement synchrony during a structured face-to-face communication task, comparing non-autistic dyads (two typically developing [TD] partners) with mixed-neurotype dyads (one TD speaker paired with one autistic listener), using gyroscope-based tracking and time-resolved trajectory modelling. Phase-based synchrony, indexed by the phase-locking value (PLV), was lower overall in mixed-neurotype dyads. Critically, time-resolved analyses revealed a marked group difference in synchrony trajectories: non-autistic dyads showed progressive, adaptive growth in synchrony over the interaction, whereas mixed-neurotype dyads showed a significantly attenuated, flatter pattern. These findings suggest that autism may involve altered temporal organization of social coordination rather than simply reduced synchrony overall.

**Lay Abstract:** When we talk with someone, we often naturally match their body language and rhythms without even realizing it. This physical “syncing up” helps us feel connected, builds trust and shared understanding, and makes communication flow easily. Research shows that autistic people might sync their movements differently during conversations compared to non-autistic people. However, past studies usually just measured an overall average of this syncing across a whole interaction. This approach misses how human interactions actually unfold over time. We wanted to know: do autistic people just sync less overall, or does their syncing change differently as the conversation goes on? To find out, we used small motion sensors to track the head movements of adults having structured face-to-face conversations and compared two types of pairs: non-autistic pairs, where both people were non-autistic, and mixed-neurotype pairs, where one non-autistic speaker talked to one autistic listener. We found a notable difference in how the two groups interacted over time. For the non-autistic pairs, the physical syncing grew progressively stronger as the conversation progressed; they progressively “tuned in” to each other. In contrast, mixed-neurotype pairs showed a flatter pattern—their level of syncing stayed relatively constant from start to finish without that same gradual build-up. These findings are important because they suggest that differences in autistic communication are not simply a “lack” or “deficit” in social coordination. Instead, autistic individuals have a distinct style of interacting—one that maintains social engagement without relying on the progressive build-up of physical syncing that non-autistic people use. Taken together, our results highlight the importance of examining how interactions evolve over time to better understand the different ways autistic and non-autistic people communicate.

## Introduction

Interpersonal synchrony broadly refers to the temporal alignment of behavior between individuals during social interaction (daSilva and Wood, 2024; Marzoratti and Evans, 2022; Schilbach and Redcay, 2025). It can be expressed across multiple modalities, including body movement, gesture, speech rhythm, and physiological activity (Cornejo et al., 2017; Delaherche et al., 2012; Dumas et al., 2010; Gordon et al., 2024; Hoehl et al., 2021). In face-to-face communication, such synchrony reflects the continuous adjustment of movement, posture, and timing between interacting partners, through which patterns of behavioral alignment emerge over time. These coordinated patterns have been associated with mutual understanding and social engagement (Gallotti et al., 2017; Oullier et al., 2008; Schmidt et al., 2011; Stolk et al., 2016), as well as with rapport, affiliation, and cooperation (Cheng et al., 2020; Hove and Risen, 2009; Lakin and Chartrand, 2003; Miles et al., 2009; Valdesolo et al., 2010). From a dynamical systems perspective, such coordination arises through reciprocal coupling between partners, leading to the spontaneous alignment of behavioral rhythms (Dumas and Fairhurst, 2021; Keller et al., 2014; Mayo and Gordon, 2020; Tognoli et al., 2020). In this sense, synchrony is understood not merely as a by-product of successful interaction, but as a core mechanism through which social behavior becomes organized and mutually intelligible (Redcay and Schilbach, 2019; Schilbach et al., 2013).

Interpersonal synchrony has attracted particular interest in autism spectrum disorder (ASD), given longstanding evidence that autistic individuals often experience difficulties in real-time social coordination (McNaughton and Redcay, 2020). A growing body of research suggests that synchrony involving autistic individuals is not necessarily absent, but is often reduced, atypical, or more variable depending on task, modality, and analytical approach (Carnevali et al., 2024; de Marchena and Eigsti, 2010; Georgescu et al., 2020; Glass and Yuill, 2024; Oberman et al., 2009). Such differences have been reported across a range of behaviors, including imitation, joint motor coordination, and naturalistic social interaction (de Marchena and Eigsti, 2010; Fitzpatrick et al., 2016; Georgescu et al., 2020; Isaksson et al., 2018; Marsh et al., 2009; Oberman et al., 2009). More broadly, this literature suggests that atypical social interaction in autism may involve altered temporal organization rather than a simple reduction in overt social behavior—a distinction important because interpersonal difficulty may lie not only in whether coordination occurs, but in the temporal pattern through which it changes during interaction.

One important limitation of the existing literature is that interpersonal synchrony has often been assessed using static or session-averaged indices (Delaherche et al., 2012; Efthimiou et al., 2025; Gordon et al., 2024; Mayo and Gordon, 2020; Ramseyer, 2020). Although these approaches have yielded important insights, they may obscure how synchrony develops over time during interaction (Gordon et al., 2024; Mayo and Gordon, 2020; Schmidt and Richardson, 2008). Social coordination is inherently dynamic: rather than being fixed from the outset, it often emerges gradually as partners become increasingly attuned to one another (Kelso, 1995; Schilbach et al., 2013; Schmidt and Richardson, 2008; Shatz et al., 2024). Recent work on the temporal dynamics of nonverbal synchrony has emphasized that clinically meaningful group differences may be expressed not only in overall synchrony magnitude, but also in the trajectory through which synchrony strengthens, stabilizes, or fails to emerge (Carnevali et al., 2024; Dumas and Fairhurst, 2021; Georgescu et al., 2020; Gordon et al., 2024; Mayo and Gordon, 2020; Ohayon and Gordon, 2025; Shatz et al., 2024). This issue may be especially important in autism, where interpersonal difficulty may lie less in the complete absence of coordination than in a reduced tendency for coordination to become progressively organized over time.

This temporal perspective is also consistent with predictive accounts of social interaction (Bastos et al., 2012; Brown and Brüne, 2012; Friston, 2005; McMahon and Isik, 2023; Tamir and Thornton, 2018). Successful coordination requires individuals not only to respond to ongoing behavior, but also to continuously update expectations about a partner’s forthcoming actions (Frith, 2012; Schurz et al., 2021; Wolpert et al., 2003). Predictive processing accounts of autism further propose that autistic individuals may be less able to use prior information flexibly to anticipate incoming sensory and social signals (Cannon et al., 2021; Lawson et al., 2014; Palmer et al., 2017; Pellicano and Burr, 2012; Sinha et al., 2014). Such differences could affect the gradual refinement of temporal expectations supporting stable coordination, such that atypical synchrony involving autistic individuals may reflect reduced efficiency in the history-sensitive processes through which interaction partners become progressively attuned to one another.

The present study examined the temporal dynamics of interpersonal synchrony during a structured face-to-face communication task, comparing non-autistic dyads (two typically developing [TD] partners) with mixed-neurotype dyads (a TD speaker paired with an autistic listener). We focused specifically on head movement, as head nods and subtle orientation changes are salient nonverbal cues in conversation and provide a tractable signal for examining temporal coordination. We adopted a unidirectional speaker–listener format to preserve stable communicative roles, reducing variability associated with rapid turn exchange and allowing temporal changes in coordination to be examined within a controlled interactional structure. To characterize how synchrony changed over the course of interaction, we applied time-resolved trajectory modeling to gyroscope-derived angular velocity signals and quantified phase-based coordination using the phase-locking value (PLV). The study asked whether interpersonal synchrony in mixed-neurotype dyads differs from that in non-autistic dyads primarily in overall phase-locking magnitude or in the temporal trajectory through which phase synchrony changes across the interaction.

## Methods

### Participants

Seventy-two adults participated, including 54 non-autistic (typically developing; TD) individuals and 18 autistic adults. Participants were grouped into two non-overlapping dyad types: 36 TD participants formed 18 non-autistic dyads (TD–TD), and the remaining 18 TD participants were each paired with one autistic participant to form 18 mixed-neurotype dyads (TD–autistic). No participant took part in more than one dyad type.

All TD participants were undergraduate students (mean age: 21.0 years, SD = 0.92). Mean age among autistic participants was 23.0 years (SD = 1.65). All participants were native Japanese speakers with normal hearing and normal or corrected-to-normal vision. To minimize effects of prior familiarity, dyads consisted of same-sex individuals who were previously unacquainted (male:female = 1:1). In mixed-neurotype dyads, all 18 autistic participants served as listeners and were paired with TD speakers; this group comprised 9 males and 9 females.

Autistic participants were required to have a prior clinical diagnosis of autism spectrum disorder according to DSM-5 criteria, established by a licensed physician and verified by official documentation. Individuals with intellectual disability, severe psychiatric comorbidity, or major neurological disease were excluded. TD participants reported no history of developmental, psychiatric, or neurological disorders. All autistic participants completed the Japanese version of the Adult Autism Spectrum Quotient (AQ). The autistic participants had a mean AQ score of 30.56 (SD = 7.47) and mean IQ/DQ of 96.88 (SD = 17.61; numeric values available for 16/18 participants). Two participants (11.1%) were currently medicated. Comorbidities were reported in six participants: ADHD (n = 4), specific learning disorder (n = 2), epilepsy (n = 1), and color vision deficiency (n = 1); one participant had both ADHD and SLD. Clinical and demographic characteristics are summarized in Table 1.

**Table 1.** Clinical and demographic characteristics of autistic participants. Values are M (SD) unless otherwise noted. All autistic participants served as listeners and were paired with a TD speaker in a mixed-neurotype dyad. Sex balance was enforced (male/female = 1:1) across dyads.

| Characteristic | Autistic participants | Notes |
| --- | --- | --- |
| Dyads, n | 18 autistic listeners (each paired 1:1 with a TD speaker in a mixed-neurotype dyad) | Total mixed-neurotype dyads = 18 |
| Sex, n (male/female) | 9 / 9 | Same-gender strangers in each dyad |
| Age (years) | 23.0 (1.65) |  |
| AQ score (Japanese Adult AQ) | 30.56 (7.47) | Autistic group range: 10–38 |
| IQ/DQ | 96.88 (17.61)* | *Numeric values for 16/18; 2/18 recorded as “average range” |
| Currently medicated, n (%) | 2 (11.1%) |  |
| Comorbidities, n | ADHD: 4; SLD: 2; Epilepsy: 1; Color-vision deficiency: 1 | Multiple entries possible (one participant had SLD+ADHD) |

This study was approved by the Ethics Committee of Kyoto University of Education (Approval No. 1805) and conducted in accordance with the Declaration of Helsinki. Written informed consent was obtained from all participants, with additional parental or guardian consent when required.

### Experimental Environment and Apparatus

The experiment was conducted in a controlled indoor setting with participants seated 1.8 m apart facing one another. Environmental conditions were kept stable throughout testing (mean room temperature: 20.7°C; illuminance: 1,432 lx; ambient noise: 31.4 dB). Head movement was recorded using wireless triaxial motion sensors with built-in gyroscopes (TSND121, ATR-Promotions, Kyoto, Japan; approx. 37 × 46 × 12 mm, 22 g) attached to participants’ foreheads using hypoallergenic tape. A third sensor (TSND151, ATR-Promotions) served as a temporal reference device for synchronization cues at the start and end of each session. Time-stamped data were transmitted via Bluetooth to a laptop at 100 Hz. The gyroscope range was set to ±2000 dps (angular velocity stored at 0.01 dps resolution). Video recording was used to confirm task compliance only.

### Experimental Procedure

A structured face-to-face communication task was used to elicit spontaneous head movements under controlled but ecologically relevant conditions. In each dyad, one participant acted as speaker and the other as listener. Speakers were given a printed text entitled Cashless Society, adapted from the Japanese-language version of Wikipedia (∼2,600 Japanese characters)—a contemporary, emotionally neutral topic. They were given preparation time but were not permitted to take notes or rehearse aloud. During the interaction, speakers explained the content in their own words in a natural speaking style, without reading directly from the text. Listeners were asked not to interrupt but were encouraged to display natural listening responses, including nodding and facial expressions indicating attention.

In non-autistic dyads, speaker–listener roles were assigned at random. In mixed-neurotype dyads, the TD participant always served as speaker and the autistic participant as listener, focusing on receptive aspects of interpersonal coordination. Task instructions for autistic listeners were provided in a simplified format with visual aids and additional time for clarification. TD speakers were not informed of their partner’s diagnostic status, minimizing expectancy effects. Recording onset and offset were marked by the experimenter’s claps. All other aspects of the procedure— room setup, task content, and recording—were identical across dyad types. Additional apparatus and procedural details are provided in Supplementary Materials S1.

### Data Analysis

#### Signal Preprocessing

To define the valid interaction period, a synchronization dataset containing explicit start/end markers was used. For each recording, tri-axial angular velocity signals from the synchronization file were converted to an angular velocity magnitude signal, and prominent peaks corresponding to the synchronization movements were identified. The interaction window was then defined using the first and last valid synchronization peaks, reflecting the intended start and end of the task. Speaker and listener recordings were subsequently cropped to this interval.

The primary signal was the magnitude of tri-axial angular velocity, calculated as the Euclidean norm of the three gyroscope axes after converting raw outputs to deg/s. Angular velocity was selected because it directly captures rotational head movements. Signals were linearly detrended to reduce slow drift and improve stationarity, then band-pass filtered using a fourth-order zero-phase Butterworth filter with cutoffs of 0.1-5 Hz to capture conversational head movement rhythms, including nodding-related oscillations, while removing slow drift and high-frequency noise (Fujiwara et al., 2020; Hale et al., 2020; Yokozuka et al., 2018). Filtering was applied in a forward-backward manner to avoid phase distortion. Filtered signals were downsampled from 100 Hz to 10 Hz by decimation, preserving information within the band of interest while reducing computational cost for time-resolved synchrony modeling (see Supplementary Materials S2 for details).

#### Phase-Based Synchrony Metrics

Because conversational head motion often exhibits low-frequency oscillatory structure, synchrony can be expressed as phase alignment between dyad members. We therefore used the phase-locking value (PLV) as the main synchrony index (Lachaux et al., 1999; Mormann et al., 2000). Instantaneous phase was extracted from the filtered signal using the Hilbert transform (Boashash, 1992), and speaker-listener phase differences were used to compute PLV. PLV ranges from 0 to 1, with higher values indicating more consistent phase alignment over time. To capture dynamic changes in synchrony, PLV was also computed within overlapping sliding windows (5 s width, 1 s step), producing a window-level trajectory (*PLV*_*w*_) for each dyad. Summary indices included mean sliding-window PLV and its standard deviation across windows, representing the strength and stability of dynamic phase synchrony, respectively. Further details are provided in Supplementary Materials S3.

#### Time Normalization and Statistical Analyses

Because interaction duration varied across dyads, window center times were normalized to [0,1] within each dyad (0 = early interaction, 1 = late interaction). Session-level metrics were compared between groups using Welch’s independent-samples t-tests, with Cohen’s d as effect size. Time-resolved synchrony measures were analyzed using linear mixed-effects models with dyad as a random intercept (Bates et al., 2015; Singer and Willett, 2003; Kenny et al., 2020). Window-level measures were summarized into 21 equally spaced normalized-time bins per dyad, and the following model was fitted:

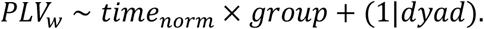

These models tested (i) a main effect of dyad type, reflecting overall synchrony differences; (ii) a main effect of time, reflecting overall temporal change; and (iii) the Time × Group interaction, reflecting group differences in PLV trajectories. As a complementary dyad-level summary of temporal change, window-level PLV values were averaged separately for the early (*time*_*norm*≤_0.5) and late (*time*_*norm*_ >0.5) halves of each interaction, and late-early differences were tested using within-group paired t-tests. To verify that observed synchrony reflected genuine interpersonal motor coupling rather than spurious overlap from shared task structure or movement autocorrelations, we conducted a permutation-based pseudo-synchrony analysis (Ramseyer and Tschacher, 2011): pseudo-dyads were generated by randomly shuffling partners within each group (5,000 derangements) while preserving communicative roles, and empirical p-values were computed by comparing observed group means with the pseudo-null distributions (see Supplementary Materials S4 for details). All analyses were performed in Python using SciPy and Statsmodels (α = .05, two-tailed).

## Results

All 36 dyads (non-autistic: n = 18; mixed-neurotype: n = 18) were successfully processed. Time-resolved analyses used 5 s sliding windows (1 s step), yielding 20,159 windows in total (mixed-neurotype: M = 594.28 per dyad; non-autistic: M = 525.67 per dyad). Window center times were normalized to [0,1] within each dyad and discretized into 21 equally spaced bins for trajectory visualization.

### Phase-Based Synchrony

#### Session-Level Phase Synchrony

Session-level phase synchrony was quantified using global PLV, summarizing the overall consistency of speaker–listener phase alignment across the full interaction. Non-autistic dyads showed higher global PLV than mixed-neurotype dyads (non-autistic: M = 0.227, SD = 0.084; mixed-neurotype: M = 0.160, SD = 0.101), a significant group difference, Welch’s t(32.90) = 2.134, p = .040, Cohen’s d = 0.711. This indicates that mixed-neurotype dyads exhibited reduced overall phase-based head-motion synchrony during unidirectional interaction.

#### Time-Resolved Phase Synchrony

Time-resolved phase synchrony was examined using sliding-window PLV. Mean sliding-window PLV was higher in non-autistic than mixed-neurotype dyads (non-autistic: M = 0.348, SD = 0.056; mixed-neurotype: M = 0.295, SD = 0.069), Welch’s t(32.54) = 2.533, p = .016, d = 0.844, indicating that group differences were more pronounced in dynamic phase synchrony indices than in session-level averages alone.

Trajectory modeling of window-level PLV (outcome ∼ timenorm × group + (1|dyad)) revealed a significant Time × Group interaction (z = 2.253, p = .024), indicating that PLV increased more strongly over time in non-autistic than in mixed-neurotype dyads. Full trajectory-model results are provided in Supplementary Materials S5. As shown in Figure 1a, the observed dyad-equal binned trajectories revealed consistently higher PLV in non-autistic than mixed-neurotype dyads and a clearer increase over normalized interaction time in non-autistic dyads. Figure 1b displays the underlying dyad-bin observations together with the corresponding mixed-effects fitted lines, illustrating that the model captured a positive temporal trend in non-autistic dyads and a comparatively flatter trajectory in mixed-neurotype dyads. Figure 1c shows the model-estimated group difference in window-level PLV across normalized time, indicating that the between-group difference widened progressively over the course of the interaction.

**Figure 1.**
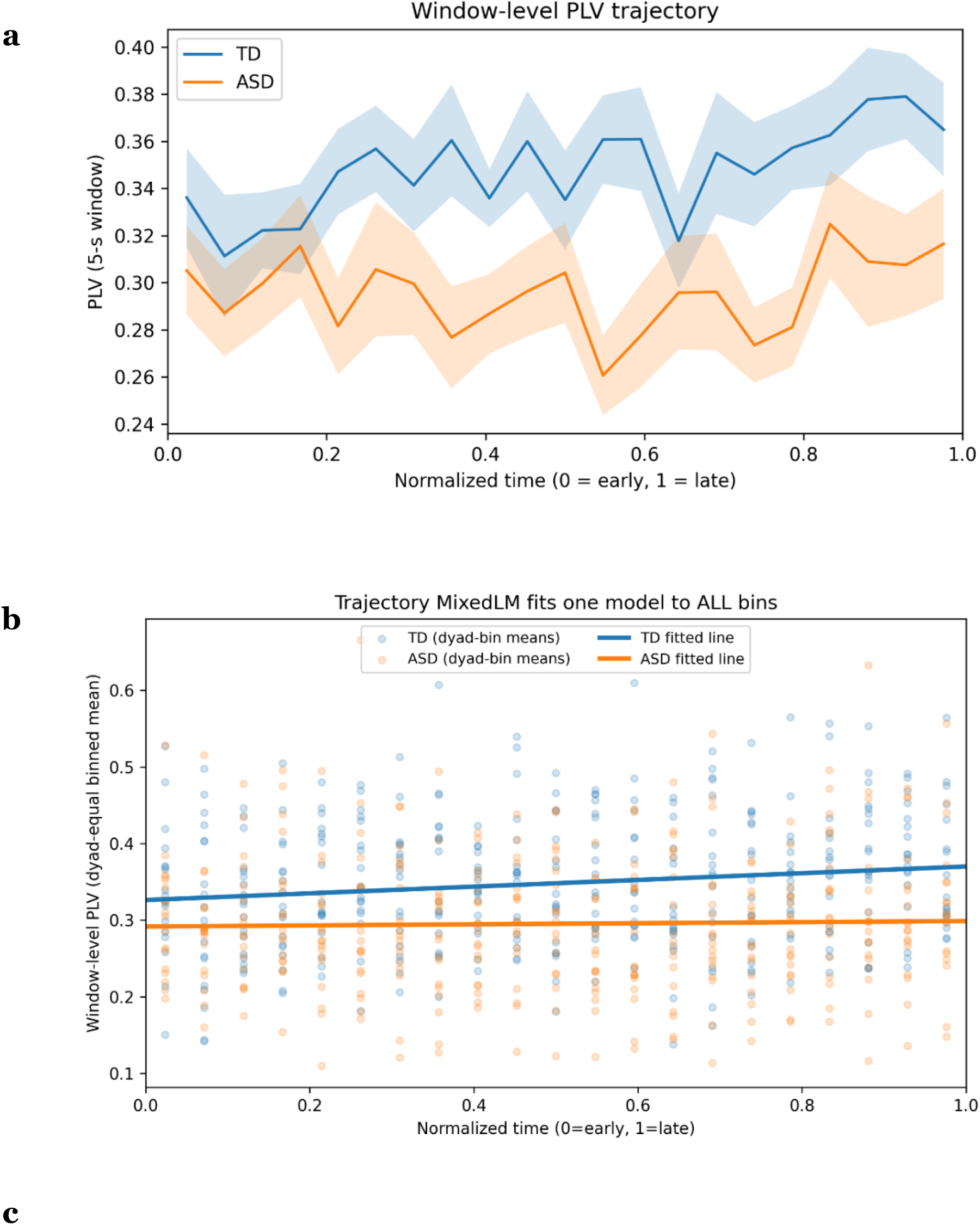
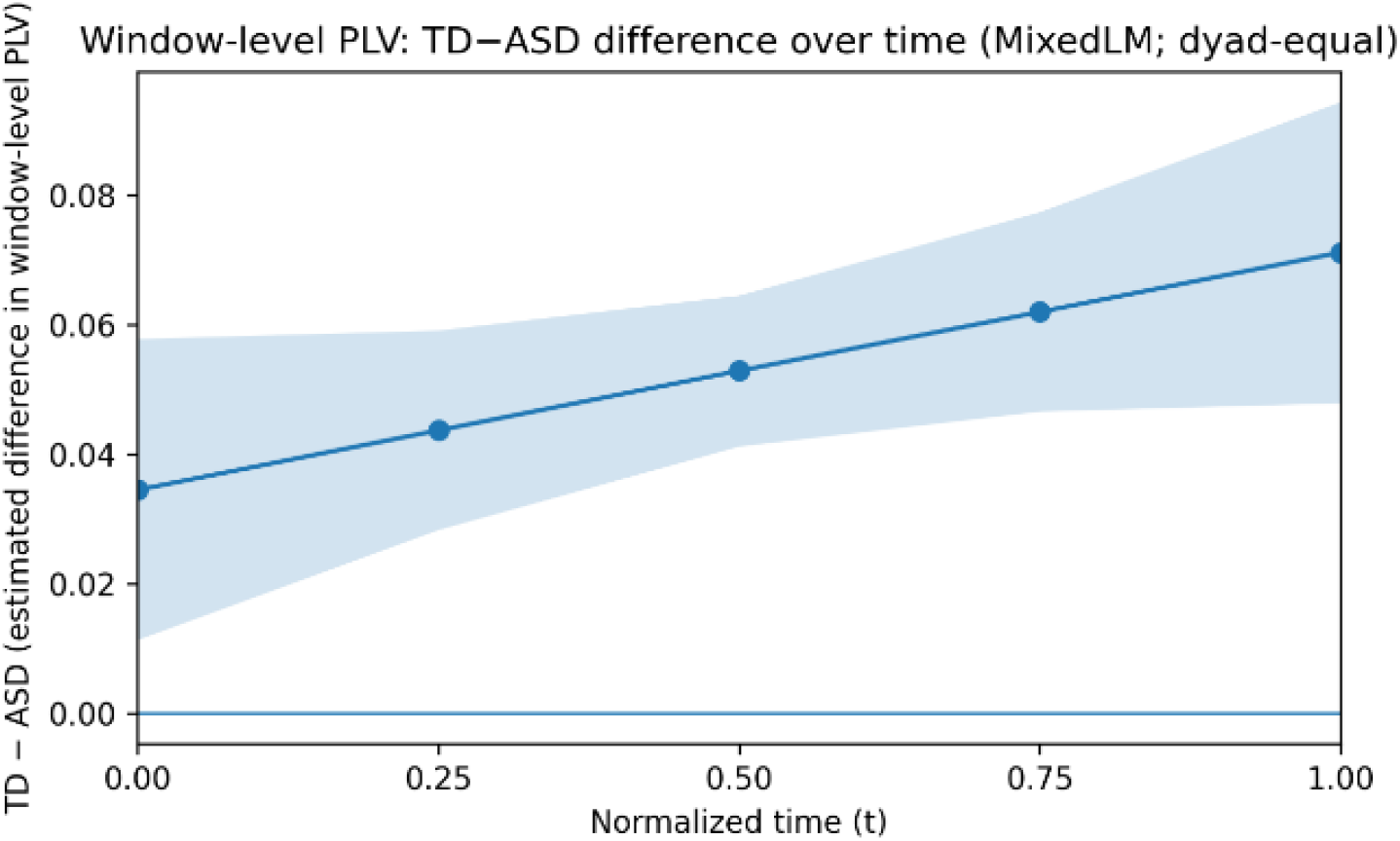
Time-resolved phase synchrony trajectories and model-estimated group differences. (a) Window-level PLV trajectories for non-autistic and mixed-neurotype dyads across normalized interaction time (0 = early, 1 = late); solid lines = dyad-equal group means across 21 bins, shaded regions = ± s.e.m. (b) Dyad-bin observations with mixed-effects fitted lines. (c) Model-estimated group difference in window-level PLV across normalized time (solid line = estimated difference, shaded region = 95% CI).

As a complementary dyad-level summary, non-autistic dyads showed significantly higher PLV in the late than early half of the interaction (mean late–early ΔPLV = +0.017; paired t(17) = 2.156, p = .046), whereas mixed-neurotype dyads showed no early–late change (ΔPLV = −0.001; paired t(17) = −0.092, p = .928). Together, these results indicate that the principal dynamic group difference lay not only in higher overall phase synchrony in non-autistic dyads, but also in a stronger time-dependent growth of synchrony—an adaptive growth trajectory that was significantly attenuated in mixed-neurotype dyads.

### Additional Analyses

Pearson correlations tested whether interaction duration was associated with phase synchrony. For mean sliding PLV versus duration, correlations were small in both groups (non-autistic: r = −0.144, p = .570; mixed-neurotype: r = −0.033, p = .895). For global PLV versus duration, correlations were likewise small (non-autistic: r = −0.071, p = .779; mixed-neurotype: r = −0.144, p = .568). These results provide no evidence that longer sessions were associated with systematic synchrony decay; the primary group difference concerned time-resolved synchrony growth.

To evaluate whether the observed PLV values reflected genuine interpersonal motor entrainment rather than spurious correlations, we compared observed synchrony against a surrogate null distribution from 5,000 pseudo-dyad permutations. Non-autistic dyads’ global PLV (M = 0.227) substantially exceeded the pseudo-null baseline (M = 0.075, SD = 0.005; Z = 29.04, p < .0002). Mixed-neurotype dyads’ global PLV (M = 0.160) likewise exceeded its pseudo-null (M = 0.062, SD = 0.006; Z = 17.89, p < .0002). Identical patterns (all p < .0002) were observed for mean sliding-window PLV (Figure 2). These findings confirm that synchrony in both groups exceeded chance.

**Figure 2.**
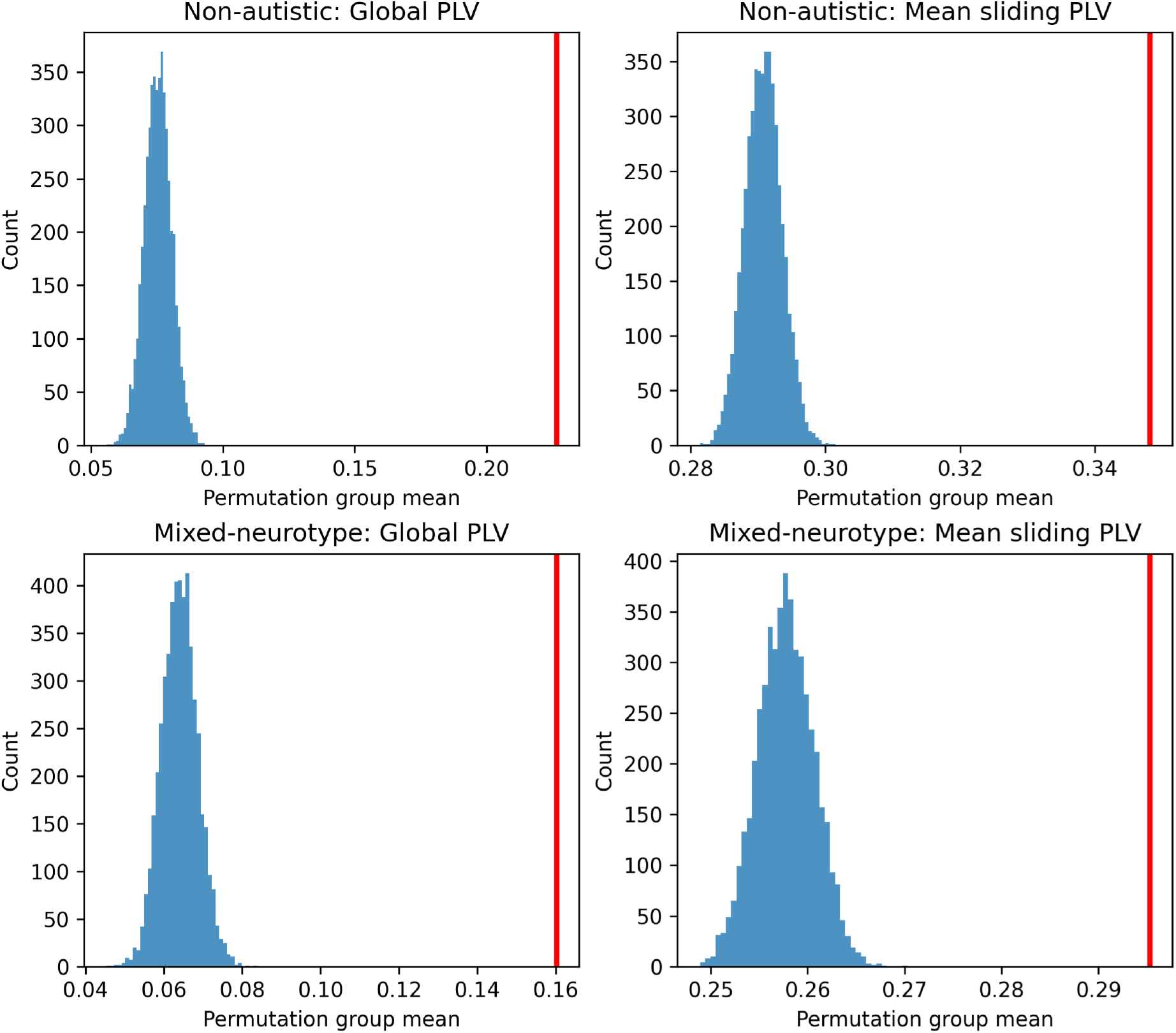
Permutation-based pseudo-synchrony distributions. Histograms show null distributions of group-mean synchrony from 5,000 pseudo-dyad permutations (partner shuffling within groups, preserving speaker–listener roles). Red vertical lines indicate observed group means. Rows: non-autistic dyads (top) and mixed-neurotype dyads (bottom). Columns: global PLV and mean sliding-window PLV.

## Discussion

The present study examined temporal dynamics of head-movement synchrony during unidirectional speaker–listener interaction in non-autistic and mixed-neurotype adult dyads. Two main findings emerged. First, mixed-neurotype dyads showed reduced phase-based synchrony at both the session level and the dynamic window level. Second, time-resolved analyses revealed a group-specific temporal pattern: non-autistic dyads showed progressive increases in phase synchrony across the interaction, whereas mixed-neurotype dyads showed attenuated growth, as reflected in the significant Time × Group interaction for PLV. Thus, the most salient group difference lay not simply in whether partners moved more similarly overall, but in whether their temporal coordination became increasingly aligned over time.

These findings align with the broader literature showing that interpersonal synchrony involving autistic individuals is often present but reduced, atypical, or more variable depending on task, modality, and analytic approach (Carnevali et al., 2024; de Marchena and Eigsti, 2010; Georgescu et al., 2020; Glass and Yuill, 2024; Oberman et al., 2009). The present study extends this literature by shifting emphasis from overall synchrony level to whether synchrony becomes progressively stronger over interaction. If group differences are expressed more clearly in the temporal emergence of synchrony than in static mean levels, session-averaged analyses may overlook an important feature of the coordination profile associated with autism. The present results suggest that mixed-neurotype dyads may be characterized not simply by reduced synchrony, but by a reduced tendency for synchrony to become increasingly organized as interaction unfolds.

### Adaptive Synchrony Growth in Non-Autistic Dyads and Its Attenuation in Mixed-Neurotype Dyads

A central implication of these findings is that non-autistic dyads exhibited adaptive growth of synchrony over the course of the interaction, whereas mixed-neurotype dyads showed a reduced tendency for such growth. The temporal increase in phase alignment observed in non-autistic dyads likely reflects a gradual process of interpersonal adjustment, whereby participants became increasingly attuned to the temporal structure of their partner’s movements (Hasson and Frith, 2016; Keller et al., 2014). This is consistent with theoretical perspectives emphasizing that successful joint action is not fixed from the outset but emerges spontaneously through ongoing integration of perception, attention, and mutual adaptation across multiple timescales (Hendrikse et al., 2023; Keller et al., 2014; Keller et al., 2026; Schmidt and Richardson, 2008; Sebanz et al., 2006). Coordination from this perspective depends on maintaining both temporal precision and flexibility, such that partners continuously correct small timing discrepancies while accommodating broader interactional demands (Gallotti et al., 2017; Gordon et al., 2024; Hoehl et al., 2021; Mayo and Gordon, 2020; Ravreby et al., 2022). The non-autistic trajectory may thus index the progressive stabilization of a dynamically coupled interpersonal system (Miles et al., 2009; Schmidt et al., 2011; Shatz et al., 2024). This does not imply that “more synchrony” is universally adaptive in all contexts (daSilva and Wood, 2024; Mayo and Gordon, 2020); rather, in the present role-structured task, the gradual increase in phase alignment likely reflects context-appropriate attunement toward a more efficient and stable interpersonal timing regime.

By contrast, the flatter trajectory in mixed-neurotype dyads suggests not the absence of coordination itself, but a reduced tendency for coordination to become progressively stabilized across the interaction. These dyads appeared to remain at a relatively constant level of coordination without showing the incremental optimization observed in non-autistic dyads, suggesting reduced evidence for the progressive build-up of dynamic interpersonal adjustments through which coordination becomes more temporally organized as interaction unfolds (Bolis et al., 2018; Dumas and Fairhurst, 2021; Hasson and Frith, 2016; Keller et al., 2014; Mayo and Gordon, 2020; Tognoli et al., 2020). More broadly, this interpretation is consistent with the view that interpersonal difficulty involving autistic individuals may reflect altered temporal organization of social coordination rather than merely a global reduction in overt social behavior (de Marchena and Eigsti, 2010; Fitzpatrick et al., 2016; Isaksson et al., 2018; Marsh et al., 2013; Ohayon and Gordon, 2025). Relatedly, recent work on natural conversation found no robust group difference in overall social motor synchrony between autistic and non-autistic dyads, but showed that synchrony was more strongly linked to rapport in non-autistic dyads, suggesting that the interpersonal significance of synchrony may differ by neurotype (Efthimiou et al., 2025). The flatter trajectory observed here may thus reflect a distinct interactional style sustained without the progressive, synchrony-based temporal alignment characteristic of non-autistic dyads, rather than disengagement per se.

### Predictive Coding and Adaptive Synchrony Growth

The increasing synchrony trajectory in non-autistic dyads is broadly consistent with predictive accounts of social interaction, which propose that successful coordination depends on the continuous formation and updating of expectations about a partner’s forthcoming actions (Brown and Brüne, 2012; Frith, 2012; Frith and Frith, 1999; Kilner et al., 2007; Koster-Hale and Saxe, 2013). Rather than arising from mere co-occurrence of movement, the non-autistic growth pattern likely reflects an adaptive process of interpersonal calibration, in which partners progressively attune to each other’s nonverbal rhythms and thereby enhance mutual predictability over time (Chartrand and Bargh, 1999; Friston and Frith, 2015; Keller et al., 2014; Marsh et al., 2009). As interaction unfolds, prior sensorimotor information may be integrated into increasingly stable expectations about the partner’s behavior, allowing synchrony to emerge as a temporally scaffolded and self-reinforcing property of dyadic exchange.

The flatter trajectory in mixed-neurotype dyads is consistent with accounts of altered social predictive coding in autism (Cannon et al., 2021; Lawson et al., 2017; Pellicano and Burr, 2012; Sinha et al., 2014). Bayesian accounts, including the hypo-priors hypothesis, propose that autistic individuals may be less able to use prior experience to constrain current perception and anticipate forthcoming sensory input (Lawson et al., 2014; Palmer et al., 2017; Pellicano and Burr, 2012; Sinha et al., 2014). In the present mixed-neurotype setting, this may have limited the dyad’s capacity to accumulate and exploit the temporal history of the exchange for predicting the partner’s movement phase, such that each moment of interaction is processed with reduced leverage from preceding context (Haker et al., 2016; Lawson et al., 2017; Palmer et al., 2017; Sevgi et al., 2020). The consequence is not simply lower synchrony, but reduced emergence of its time-dependent amplification, suggesting that attenuated adaptive synchrony in mixed-neurotype dyads may reflect differences in the predictive, history-sensitive computations through which coordination ordinarily becomes stronger over time.

### Translational Implications and Limitations

A key interpretive question is whether reduced synchrony in mixed-neurotype dyads reflects progressive disengagement over time. The present findings do not support a simple monotonic decline with longer interaction; duration–synchrony correlations were negligible in both groups, and surrogate data analyses confirmed that synchrony remained above chance, indicating attenuated adaptive attunement rather than a complete absence of coordination. Instead, the principal group difference was the attenuated emergence of adaptive growth—a clinically meaningful distinction suggesting that interpersonal coordination in mixed-neurotype interactions may remain functionally engaged yet fail to converge toward increasingly efficient temporal attunement. In practice, this may manifest as interactions that feel stable but do not become progressively more aligned over time. Because the autistic participants were adults without intellectual disability, these findings further suggest that reduced adaptive synchronization is not readily explained by generalized cognitive impairment, but may reflect a more specific difference in interpersonal timing processes (Masi et al., 2017; Matson and Shoemaker, 2009). This distinction is also clinically relevant: rather than focusing exclusively on increasing overt social behaviors, interventions might also support the micro-temporal adjustment processes through which interpersonal timing becomes more finely tuned during ongoing interaction (Bolis et al., 2018; McNaughton and Redcay, 2020). Consistent with prior work on phase differences in body movement (Kwon et al., 2015; Kwon & Kotani, 2025), sliding-window PLV trajectory analysis may offer a promising quantitative framework for tracking such changes and for evaluating intervention outcomes across different contexts.

Several limitations should guide future research. First, while gyroscope-based head motion provides high temporal resolution and direct access to rotational movement, it does not capture facial affect, gaze, or speech prosody; multimodal integration may better explain how synchrony emerges in conversation (Gordon et al., 2024; Ohayon & Gordon, 2025). Second, the unidirectional design is both a strength, because it stabilizes speaker–listener roles, and a constraint, because it limits opportunities for mutual adaptation during reciprocal interaction (Kwon, 2025; Kwon & Kotani, 2023). Extending this framework to bidirectional conversation, autistic-speaker conditions, and autistic-autistic dyads would help clarify whether dyad-type differences reflect differences in role adaptation, responsiveness to cues, predictive timing mechanisms, or broader features of cross-neurotype interaction. Finally, given the heterogeneity of autism, future work should examine whether adaptive synchrony trajectories vary as a function of autism phenotype, symptom severity, social motivation, and communicative ability.

## Conclusion

In adult unidirectional speaker–listener interaction, non-autistic dyads demonstrated adaptive growth of phase-based head-motion synchrony over time, whereas mixed-neurotype dyads showed attenuated emergence of this adaptive alignment. This dynamic signature was most clearly captured by time-resolved phase synchrony (PLV). These findings highlight the importance of analyzing how synchrony evolves during interaction and suggest that reduced adaptive synchronization—rather than simple synchrony loss or progressive disengagement—may be a key characteristic of interpersonal coordination in mixed-neurotype adult dyads involving autistic individuals without intellectual disability.

## Acknowledgements

The authors thank all participants who took part in this study.

## Author contributions

J.K. and H.K. conceived and designed the study. J.K. and H.K. performed the experiments and analysed the data. J.K. wrote the first draft of the manuscript. All authors revised and approved the final manuscript.

## References

Bastos, A. M., Usrey, W. M., Adams, R. A., Mangun, G. R., Fries, P., & Friston, K. J. (2012). Canonical microcircuits for predictive coding. Neuron, 76(4), 695–711.

Bates, D., Mächler, M., Bolker, B., & Walker, S. (2015). Fitting linear mixed-effects models using lme4. Journal of Statistical Software, 67, 1–48.

Boashash, B. (1992). Estimating and interpreting the instantaneous frequency of a signal. I. Fundamentals. Proceedings of the IEEE, 80(4), 520–538.

Bolis, D., Balsters, J., Wenderoth, N., Becchio, C., & Schilbach, L. (2018). Beyond autism: Introducing the dialectical misattunement hypothesis and a Bayesian account of intersubjectivity. Psychopathology, 50(6), 355–372.

Brown, E. C., & Brüne, M. (2012). The role of prediction in social neuroscience. Frontiers in Human Neuroscience, 6, Article 147.

Cannon, J., O’Brien, A. M., Bungert, L., & Sinha, P. (2021). Prediction in autism spectrum disorder: A systematic review of empirical evidence. Autism Research, 14(4), 604–630.

Carnevali, L., Valori, I., Mason, G., Altoè, G., & Farroni, T. (2024). Interpersonal motor synchrony in autism: A systematic review and meta-analysis. Frontiers in Psychiatry, 15, Article 1355068.

Chartrand, T. L., & Bargh, J. A. (1999). The chameleon effect: The perception–behavior link and social interaction. Journal of Personality and Social Psychology, 76(6), 893–910.

Cheng, M., Kato, M., Saunders, J. A., & Tseng, C. H. (2020). Paired walkers with better first impression synchronize better. PLoS ONE, 15(1), e0229658.

Cornejo, C., Cuadros, Z., Morales, R., & Paredes, J. (2017). Interpersonal coordination: Methods, achievements, and challenges. Frontiers in Psychology, 8, Article 1685.

daSilva, E. B., & Wood, A. (2024). How and why people synchronize: An integrated perspective. Personality and Social Psychology Review. Advance online publication.

de Marchena, A., & Eigsti, I. M. (2010). Conversational gestures in autism spectrum disorders: Asynchrony but not decreased frequency. Autism Research, 3(6), 311–322.

Delaherche, E., Chetouani, M., Mahdhaoui, A., Saint-Georges, C., Viaux, S., & Cohen, D. (2012). Interpersonal synchrony: A survey of evaluation methods across disciplines. IEEE Transactions on Affective Computing, 3(3), 349–365.

Dumas, G., & Fairhurst, M. T. (2021). Reciprocity and alignment: Quantifying coupling in dynamic interactions. Royal Society Open Science, 8(5), Article 210138.

Dumas, G., Nadel, J., Soussignan, R., Martinerie, J., & Garnero, L. (2010). Inter-brain synchronization during social interaction. PLoS ONE, 5(8), e12166.

Efthimiou, T. N., Wilks, C. E., Foster, S., Dodd, M., Sasson, N. J., Ropar, D., Lages, M., Fletcher-Watson, S., & Crompton, C. J. (2025). Social motor synchrony and interactive rapport in autistic, non-autistic, and mixed-neurotype dyads. Autism. Advance online publication.

Fitzpatrick, P., Frazier, J. A., Cochran, D. M., Mitchell, T., Coleman, C., & Schmidt, R. C. (2016). Impairments of social motor synchrony evident in autism spectrum disorder. Frontiers in Psychology, 7, Article 1323.

Friston, K. (2005). A theory of cortical responses. Philosophical Transactions of the Royal Society B: Biological Sciences, 360(1456), 815–836.

Friston, K. J., & Frith, C. D. (2015). Active inference, communication and hermeneutics. Cortex, 68, 129–143.

Frith, C. D. (2012). The role of metacognition in human social interactions. Philosophical Transactions of the Royal Society B: Biological Sciences, 367(1599), 2213–2223.

Frith, C. D., & Frith, U. (1999). Interacting minds—a biological basis. Science, 286(5445), 1692– 1695.

Fujiwara, K., Kimura, M., & Daibo, I. (2020). Rhythmic features of movement synchrony for bonding individuals in dyadic interaction. Journal of Nonverbal Behavior, 44(2), 173– 193.

Gallotti, M., Fairhurst, M. T., & Frith, C. D. (2017). Alignment in social interactions. Consciousness and Cognition, 48, 253–261.

Georgescu, A. L., Koeroglu, S., Hamilton, A. F. D. C., Vogeley, K., Falter-Wagner, C. M., & Tschacher, W. (2020). Reduced nonverbal interpersonal synchrony in autism spectrum disorder independent of partner diagnosis: A motion energy study. Molecular Autism, 11, Article 11.

Glass, D., & Yuill, N. (2024). Social motor synchrony in autism spectrum conditions: A systematic review. Autism, 28(7), 1638–1653.

Gordon, I., Tomashin, A., & Mayo, O. (2024). A theory of flexible multimodal synchrony. Psychological Review. Advance online publication.

Haker, H., Schneebeli, M., & Stephan, K. E. (2016). Can Bayesian theories of autism spectrum disorder help improve clinical practice? Frontiers in Psychiatry, 7, Article 107.

Hale, J., Ward, J. A., Buccheri, F., Oliver, D., & Hamilton, A. F. D. C. (2020). Are you on my wavelength? Interpersonal coordination in dyadic conversations. Journal of Nonverbal Behavior, 44(1), 63–83.

Hasson, U., & Frith, C. D. (2016). Mirroring and beyond: Coupled dynamics as a generalized framework for modelling social interactions. Philosophical Transactions of the Royal Society B: Biological Sciences, 371(1693), Article 20150366.

Hendrikse, S. C., Treur, J., & Koole, S. L. (2023). Modeling emerging interpersonal synchrony and its related adaptive short-term affiliation and long-term bonding: A second-order multi-adaptive neural agent model. International Journal of Neural Systems, 33(12), Article 2350038.

Hoehl, S., Fairhurst, M., & Schirmer, A. (2021). Interactional synchrony: Signals, mechanisms and benefits. Social Cognitive and Affective Neuroscience, 16(1–2), 5–18.

Hove, M. J., & Risen, J. L. (2009). It’s all in the timing: Interpersonal synchrony increases affiliation. Social Cognition, 27(6), 949–960.

Isaksson, S., Salomäki, S., Tuominen, J., Arstila, V., Falter-Wagner, C. M., & Noreika, V. (2018). Is there a generalized timing impairment in autism spectrum disorders across time scales and paradigms? Journal of Psychiatric Research, 99, 111–121.

Keller, P. E., Lee, J., Mills, P., & Harry, B. B. (2026). Modeling individual differences in temporal adaptation, anticipation, and attention allocation during rhythmic interpersonal coordination. Timing & Time Perception, 1–77.

Keller, P. E., Novembre, G., & Hove, M. J. (2014). Rhythm in joint action: Psychological and neurophysiological mechanisms for real-time interpersonal coordination. Philosophical Transactions of the Royal Society B: Biological Sciences, 369(1658), Article 20130394.

Kelso, J. A. S. (1995). Dynamic patterns: The self-organization of brain and behavior. MIT Press.

Kenny, D. A., Kashy, D. A., & Cook, W. L. (2020). Dyadic data analysis. Guilford Press.

Kilner, J. M., Friston, K. J., & Frith, C. D. (2007). Predictive coding: An account of the mirror neuron system. Cognitive Processing, 8(3), 159–166.

Koster-Hale, J., & Saxe, R. (2013). Theory of mind: A neural prediction problem. Neuron, 79(5), 836–848.

Kwon, J. (2025). Synchrony Vision: Capturing body motion synchrony through phase difference using the Kinect. IEEE Access, 13, 41658–41669.

Kwon, J., & Kotani, H. (2023). Head motion synchrony in unidirectional and bidirectional verbal communication. PLoS ONE, 18(5), Article e0286098.

Kwon, J., & Kotani, H. (2025). Quantifying body motion synchrony in autism spectrum disorder using a phase difference detection algorithm: Toward a novel behavioral biomarker. Diagnostics, 15(10), Article 1268.

Kwon, J., Ogawa, K.-I., Ono, E., & Miyake, Y. (2015). Detection of nonverbal synchronization through phase difference in human communication. PLoS ONE, 10(7), Article e0133881.

Lachaux, J.-P., Rodriguez, E., Martinerie, J., & Varela, F. J. (1999). Measuring phase synchrony in brain signals. Human Brain Mapping, 8(4), 194–208.

Lakin, J. L., & Chartrand, T. L. (2003). Using nonconscious behavioral mimicry to create affiliation and rapport. Psychological Science, 14(4), 334–339.

Lawson, R. P., Mathys, C., & Rees, G. (2017). Adults with autism overestimate the volatility of the sensory environment. Nature Neuroscience, 20(9), 1293–1299.

Lawson, R. P., Rees, G., & Friston, K. J. (2014). An aberrant precision account of autism. Frontiers in Human Neuroscience, 8, Article 302.

Lombardo, M. V., Lai, M. C., & Baron-Cohen, S. (2019). Big data approaches to decomposing heterogeneity across the autism spectrum. Molecular Psychiatry, 24(10), 1435–1450.

Marsh, K. L., Isenhower, R. W., Richardson, M. J., Helt, M., Verbalis, A. D., Schmidt, R. C., & Fein, D. (2013). Autism and social disconnection in interpersonal rocking. Frontiers in Integrative Neuroscience, 7, Article 4.

Marsh, K. L., Richardson, M. J., & Schmidt, R. C. (2009). Social connection through joint action and interpersonal coordination. Topics in Cognitive Science, 1(2), 320–339.

Marzoratti, A., & Evans, T. M. (2022). Measurement of interpersonal physiological synchrony in dyads: A review of timing parameters used in the literature. Cognitive, Affective, & Behavioral Neuroscience, 22(6), 1215–1230.

Masi, A., DeMayo, M. M., Glozier, N., & Guastella, A. J. (2017). An overview of autism spectrum disorder, heterogeneity and treatment options. Neuroscience Bulletin, 33(2), 183–193.

Matson, J. L., & Shoemaker, M. (2009). Intellectual disability and its relationship to autism spectrum disorders. Research in Developmental Disabilities, 30(6), 1107–1114.

Mayo, O., & Gordon, I. (2020). In and out of synchrony—Behavioral and physiological dynamics of dyadic interpersonal coordination. Psychophysiology, 57(6), e13574.

McMahon, E., & Isik, L. (2023). Seeing social interactions. Trends in Cognitive Sciences, 27(12), 1165–1179.

McNaughton, K. A., & Redcay, E. (2020). Interpersonal synchrony in autism. Current Psychiatry Reports, 22(3), Article 12.

Miles, L. K., Nind, L. K., & Macrae, C. N. (2009). The rhythm of rapport: Interpersonal synchrony and social perception. Journal of Experimental Social Psychology, 45(3), 585–589.

Mormann, F., Lehnertz, K., David, P., & Elger, C. E. (2000). Mean phase coherence as a measure for phase synchronization and its application to the EEG of epilepsy patients. Physica D: Nonlinear Phenomena, 144(3–4), 358–369.

Oberman, L. M., Winkielman, P., & Ramachandran, V. S. (2009). Slow echo: Facial EMG evidence for the delay of spontaneous, but not voluntary, emotional mimicry in children with autism spectrum disorders. Developmental Science, 12(4), 510–520.

Ohayon, S., & Gordon, I. (2025). Multimodal interpersonal synchrony: Systematic review and meta-analysis. Behavioural Brain Research, 480, Article 115369.

Oullier, O., De Guzman, G. C., Jantzen, K. J., Lagarde, J., & Kelso, J. A. S. (2008). Social coordination dynamics: Measuring human bonding. Social Neuroscience, 3(2), 178–192.

Palmer, C. J., Lawson, R. P., & Hohwy, J. (2017). Bayesian approaches to autism: Towards volatility, action, and behavior. Psychological Bulletin, 143(5), 521–542.

Pellicano, E., & Burr, D. (2012). When the world becomes ‘too real’: A Bayesian explanation of autistic perception. Trends in Cognitive Sciences, 16(10), 504–510.

Ramseyer, F. (2020). Motion energy analysis (MEA): A primer on the assessment of motion from video. Journal of Counseling Psychology, 67(5), 536–549.

Ramseyer, F., & Tschacher, W. (2011). Nonverbal synchrony in psychotherapy: Coordinated body movement reflects relationship quality and outcome. Journal of Consulting and Clinical Psychology, 79(3), 284–295.

Ravreby, I., Shilat, Y., & Yeshurun, Y. L. (2022). Liking as a balance between synchronization, complexity and novelty. Scientific Reports, 12, Article 3181.

Redcay, E., & Schilbach, L. (2019). Using second-person neuroscience to elucidate the mechanisms of social interaction. Nature Reviews Neuroscience, 20(8), 495–505.

Schilbach, L., & Redcay, E. (2025). Synchrony across brains. Annual Review of Psychology, 76, 883–911.

Schilbach, L., Timmermans, B., Reddy, V., Costall, A., Bente, G., Schlicht, T., & Vogeley, K. (2013). Toward a second-person neuroscience. Behavioral and Brain Sciences, 36(4), 393–414.

Schmidt, R. C., Fitzpatrick, P., Caron, R., & Mergeche, J. (2011). Understanding social motor coordination. Human Movement Science, 30(5), 834–845.

Schmidt, R. C., & Richardson, M. J. (2008). Dynamics of interpersonal coordination. In A. Fuchs & V. K. Jirsa (Eds.), Coordination: Neural, behavioral and social dynamics (pp. 281–308). Springer.

Schurz, M., Radua, J., Tholen, M. G., Maliske, L., Margulies, D. S., Mars, R. B., Sallet, J., & Kanske, P. (2021). Toward a hierarchical model of social cognition: A neuroimaging meta-analysis and integrative review of empathy and theory of mind. Psychological Bulletin, 147(3), 293–327.

Sebanz, N., Bekkering, H., & Knoblich, G. (2006). Joint action: Bodies and minds moving together. Trends in Cognitive Sciences, 10(2), 70–76.

Sevgi, M., Diaconescu, A. O., Henco, L., Tittgemeyer, M., & Schilbach, L. (2020). Social Bayes: Using Bayesian modeling to study autistic trait–related differences in social cognition. Biological Psychiatry, 87(2), 185–193.

Shatz, H., Asher, M., & Aderka, I. M. (2024). Catching a (sine) wave: Temporal dynamics of nonverbal synchrony in social anxiety disorder. Journal of Anxiety Disorders, 102, Article 102828.

Singer, J. D., & Willett, J. B. (2003). Applied longitudinal data analysis: Modeling change and event occurrence. Oxford University Press.

Sinha, P., Kjelgaard, M. M., Gandhi, T. K., Tsourides, K., Cardinaux, A. L., Pantazis, D., Diamond, S. P., & Held, R. M. (2014). Autism as a disorder of prediction. Proceedings of the National Academy of Sciences of the United States of America, 111(42), 15220–15225.

Stolk, A., Verhagen, L., & Toni, I. (2016). Conceptual alignment: How brains achieve mutual understanding. Trends in Cognitive Sciences, 20(3), 180–191.

Tamir, D. I., & Thornton, M. A. (2018). Modeling the predictive social mind. Trends in Cognitive Sciences, 22(3), 201–212.

Tognoli, E., Zhang, M., Fuchs, A., Beetle, C., & Kelso, J. A. S. (2020). Coordination dynamics: A foundation for understanding social behavior. Frontiers in Human Neuroscience, 14, Article 317.

Valdesolo, P., Ouyang, J., & DeSteno, D. (2010). The rhythm of joint action: Synchrony promotes cooperative ability. Journal of Experimental Social Psychology, 46(4), 693–695.

Wolpert, D. M., Doya, K., & Kawato, M. A. (2003). A unifying computational framework for motor control and social interaction. Philosophical Transactions of the Royal Society of London. Series B: Biological Sciences, 358(1431), 593–602.

Yokozuka, T., Ono, E., Inoue, Y., Ogawa, K.-I., & Miyake, Y. (2018). The relationship between head motion synchronization and empathy in unidirectional face-to-face communication. Frontiers in Psychology, 9, Article 1622.

