## Supplementary Materials for "Attenuated adaptive growth of interpersonal synchrony in autism"

### S1. Experimental environment, apparatus, and detailed procedure

#### Experimental Environment and Apparatus

The experiment was conducted in a controlled indoor setting in which two participants sat facing one another across a table at a distance of 1.8 m. Participants were asked not to wear hats, masks, or other head coverings. Environmental conditions were kept stable throughout testing. Mean room temperature was 20.7 °C, illuminance was 1,432 lx (CANA-0010, Tokyo Photoelectric Co., Tokyo, Japan), and ambient noise level was 31.4 dB (CHE-SD1, Sanwa Supply, Okayama, Japan).

Head movement was recorded using three wireless triaxial motion sensors with built-in gyroscopes (approx. 37 mm × 46 mm × 12 mm, 22 g). Two sensors (TSND121, ATR-Promotions, Kyoto, Japan) were attached to the participants' foreheads using hypoallergenic tape, and a third sensor (TSND151, ATR-Promotions, Kyoto, Japan) served as a temporal reference device for synchronization cues at the start and end of each session. Time-stamped data were transmitted via Bluetooth to a laptop computer at a sampling frequency of 100 Hz. The present analyses focused on gyroscope-derived angular velocity signals recorded on the X, Y, and Z axes. The gyroscope range was set to ±2000 dps, and angular velocity values were stored with a resolution of 0.01 dps. These devices also support inter-sensor time synchronization, allowing temporally aligned recordings across multiple sensors. Signal acquisition settings were configured in advance according to the manufacturer's instructions.

In addition, the interaction was video recorded using a digital camcorder (HDC-TM45, Panasonic, Osaka, Japan). These recordings were used only to confirm task compliance and general participant behaviour during the session; they were not analyzed quantitatively. Video data were anonymized and securely stored.

#### Experimental Procedure

A structured face-to-face communication task was used to elicit spontaneous head movements under controlled but ecologically relevant conditions. In each dyad, one participant acted as the speaker and the other as the listener. The interaction took place in a quiet room with stable lighting and minimal distraction. Participants sat 1.8 m apart and were instructed to maintain a natural posture and to look at their partner during the task. The paradigm was designed to resemble a natural explanatory interaction while maintaining experimental control over role structure and task content.

#### Procedure for non-autistic dyads

In non-autistic dyads, each dyad consisted of two typically developing (TD) participants, and speaker–listener roles were assigned at random. The speaker was given a printed text entitled Cashless Society, adapted from the Japanese-language version of Wikipedia. The passage contained approximately 2,600 Japanese characters and was selected because it addressed a contemporary but emotionally neutral topic that was unlikely to evoke strong personal involvement or substantial prior knowledge. Speakers were allowed time to read the material and prepare mentally before the task, but they were not permitted to take notes or rehearse aloud.

During the interaction, speakers were instructed to explain the content of the text in their own words, using a natural speaking style and typical eye contact. They were asked not to read directly from the text and to avoid exaggerated bodily movements. Listeners were instructed not to interrupt or ask questions. At the same time, they were encouraged to display natural listening responses, including nodding, short acknowledgments and facial expressions indicating attention or understanding. Participants were also instructed not to touch the sensors or change their seating position during recording.

At the start of the session, the experimenter stood at the side of the room, announced the beginning of the experiment, and produced a single clap. This served as a temporal marker recorded by the synchronization sensor and the video camera. The experimenter then left the room to minimize reactivity. When the speaker had finished the explanation, they rang a small bell placed on the table. The experimenter returned, announced the end of the session, and delivered a second clap to mark the offset of the recording. These markers were later used to define the active analysis interval.

Throughout the session, a trained researcher monitored participants for signs of fatigue or discomfort. Monitoring took place remotely via live audio–video feed from an adjacent room so that safety could be maintained without disturbing the natural flow of the interaction.

#### Procedure for mixed-neurotype dyads

In mixed-neurotype dyads, each dyad consisted of one TD speaker and one autistic listener. By design, the TD participant always served as the speaker and the autistic participant always served as the listener. This fixed-role arrangement was adopted to focus specifically on receptive aspects of interpersonal coordination, particularly the extent to which autistic listeners aligned with the speaker’s ongoing movement timing and communicative rhythm.

All other aspects of the procedure were matched to those of non-autistic dyads. The non-autistic speaker received the same Cashless Society passage and followed the same preparation and speaking instructions. Autistic listeners were given the task instructions in a simplified and supportive format, with visual aids and additional verbal explanation when necessary to ensure full understanding. Apart from these accommodations, task content, room setup, and recording procedures were identical across dyad types.

To reduce expectancy effects, non-autistic speakers were not informed of their partner’s diagnostic status. Participants were told only that they would take part in a communication task with another person. As a result, speakers in both dyad types were effectively blind to whether their partner was autistic or non-autistic. This procedure was intended to minimize the possibility that speaker behaviour would be influenced by assumptions about the listener.

The task was designed to permit the spontaneous emergence of head-movement synchrony while maintaining consistency in communicative direction, task demands, and environmental context across dyads. This allowed direct comparison of interpersonal movement coordination between dyad types.

### S2. Signal preprocessing details

The primary signal for synchrony analyses was derived from the tri-axial angular velocity (gyroscope) channels. Angular velocity was selected because it directly captures rotational head movements associated with conversational nodding and subtle head orientation adjustments, and it is less affected than acceleration by gravitational components and translational artifacts (Cook, 2016). The gyroscope measurement range was set to ±2000 dps. Raw gyroscope outputs were recorded in 0.01 dps units and were therefore converted to deg/s prior to magnitude computation and subsequent preprocessing. For each participant, angular velocity magnitude was computed as:

$$\omega(t) = \sqrt{\omega_x(t)^2 + \omega_y(t)^2 + \omega_z(t)^2},$$

where $\omega_x(t)$, $\omega_y(t)$, and $\omega_z(t)$ denote the angular velocity signals along the three axes (in deg/s). This magnitude representation provided an orientation-independent index of rotational head movement intensity suitable for dyadic synchrony analysis.

### S3. Phase-based synchrony metrics

**Phase Extraction and Phase-Based Synchrony Metrics**

Following preprocessing, instantaneous phase was extracted from the filtered angular velocity magnitude signal using the Hilbert transform. Specifically, for each participant's time series $\omega(t)$, the analytic signal $z(t)$ was computed as:

$$z(t) = \omega(t) + i\mathcal{H}[\omega(t)],$$

where $\mathcal{H}[\cdot]$ denotes the Hilbert transform. Instantaneous phase was then defined as:

$$\phi(t) = \arg(z(t)).$$

For each dyad, the instantaneous phase difference between the speaker and listener was computed as:

$$\Delta\phi(t) = \phi_{\text{speaker}}(t) - \phi_{\text{listener}}(t),$$

and wrapped to the principal interval $[-\pi, \pi]$ to preserve circular properties. The phase-difference time series $\Delta\phi(t)$ served as the basis for quantifying both session-level and time-resolved phase synchrony.

**Global Phase-Locking Value (Session-Level PLV)**

Overall phase synchrony during the interaction was quantified by the phase-locking value (PLV) computed across the entire session:

$$PLV = \left| \frac{1}{N} \sum_{t=1}^{N} e^{i\Delta\phi(t)} \right|.$$

PLV ranges from 0 to 1, with higher values indicating more consistent phase alignment across time.

**Sliding-Window PLV (Time-Resolved PLV)**

To capture dynamic changes in synchrony over the course of the interaction, PLV was computed within overlapping sliding windows. Each window had a duration of 5 seconds, with a step size of 1 second. This produced a time series of window-level PLV values ($PLV_w$) for each dyad:

$$PLV_w(k) = \left| \frac{1}{N_w} \sum_{t \in \text{window } k} e^{i\Delta\phi(t)} \right|,$$

where $N_w$ denotes the number of samples in the $k$-th window. From these window-level values, summary indices were computed including the mean sliding PLV (average of $PLV_w$) and the variability of sliding PLV (standard deviation across windows), representing the strength and stability of dynamic phase synchrony, respectively. Window-level PLV was used to quantify (i) the strength of local phase alignment at each time point and (ii) how this alignment changed as the interaction progressed.

### S4. Pseudo-Synchrony Analysis

To verify that the observed phase synchrony represents genuine interpersonal motor coupling rather than spurious overlap arising from shared experimental task structure or intrinsic movement autocorrelations, we conducted a robust permutation-based surrogate data analysis (Ramseyer and Tschacher, 2011). Pseudo-dyads were generated by randomly shuffling partners within each respective group while strictly preserving their original communicative roles. Specifically, a time series from a speaker in one dyad was paired with a time series from a listener in a different dyad within the same group, ensuring that original true pairings were explicitly excluded (i.e., a complete derangement).

For each generated pseudo-dyad, the identical preprocessing and synchrony quantification pipeline used in the main analysis—including detrending, band-pass filtering, Hilbert transformation, and the 5-second sliding window PLV calculation—was strictly applied.

We established empirical null distributions for the phase-based synchrony metrics (global PLV and mean sliding-window PLV) by performing 5,000 repeated one-to-one derangement permutations. To quantify the magnitude of genuine phase synchrony relative to chance level, we computed the effect size (Z-score) by comparing the observed group mean against the mean and standard deviation of the pseudo-null distribution. Finally, empirical p-values were calculated as the proportion of pseudo-group means that were equal to or greater than the observed actual group mean.

**S5. Mixed-effects trajectory model for time-resolved phase synchrony (window-level PLV). ($p < 0.05$: *, $p < 0.01$: **, $p < 0.001$: ***)**

| Model | Term | Estimate | SE | Stat | p |
|---|---|---|---|---|---|
| $PLV_w$ | Intercept | 0.2918 | 0.0158 | 18.416 | < 0.001*** |
| | Dyad type (non-autistic vs mixed-neurotype) | 0.0346 | 0.0224 | 1.543 | 0.1229 |
| | Time (normalized) | 0.0072 | 0.0115 | 0.622 | 0.5342 |
| | Time × Group | 0.0367 | 0.0163 | 2.253 | 0.0242* |
